# Enabling high throughput target-based drug discovery in 3D cell cultures through novel bioprinting workflow

**DOI:** 10.1101/2021.04.21.440768

**Authors:** Martin Engel, Lisa Belfiore, Behnaz Aghaei, Margareta Sutija

## Abstract

Advanced three-dimensional cell culture techniques have been adopted in many laboratories to better model *in vivo* tissue by recapitulating multi-cellular architecture and the presence of extracellular matrix features. We describe here a 3D cell culture platform in a small molecule screening workflow that uses traditional biomarker and intracellular kinase end point assay readouts. By combining the high throughput bioprinter Rastrum with the high throughput screening assay AlphaLISA, we demonstrate the utility of the workflow in 3D synthetic hydrogel cultures with breast cancer (MDA-MB-231 and MCF-7) and fibroblast cells. To establish and validate the workflow, we treated the breast cancer cultures with doxorubicin, while fibroblast cultures were stimulated with the pro-inflammatory lipopolysaccharide. 3D and 2D MDA-MB-231 cultures were equally susceptible to doxorubicin treatment, while showing opposite ERK phosphorylation changes. Doxorubicin readily entered embedded MCF-7 spheroids and markedly reduced intracellular GSK3β phosphorylation. Furthermore, quantifying extracellular interleukin 6 levels showed a very similar activation profile for fibroblasts in 2D and 3D cultures, with 3D fibroblast networks being more resistant against the immune challenge. Through these validation experiments we demonstrate the full compatibility of the bioprinted 3D cell cultures with several widely-used 2D culture assays. The efficiency of the workflow, minimal culture handling, and applicability of traditional screening assays, demonstrates that advanced encapsulated 3D cell cultures can be used in 2D cell culture screening workflows, while providing a more holistic view on cell biology to increase the predictability to *in vivo* drug response.

## Introduction

In the context of high throughput screening (HTS) for biological research, 2D cell cultures are the widespread model of choice for *in vitro* testing of novel compounds ^1^. This is in large part due to the existence of countless highly optimised workflows for the generation and analysis of 2D cell culture models, making the use of 2D cell cultures practical and cost-effective in HTS ^2^. However, growing evidence indicates that cells cultured in 2D do not sufficiently model the complex biology of *in vivo* tissues to reliably predict *in vivo* drug responses ^3^. This fundamental limitation of 2D cell culture models arises primarily from lacking the 3D tissue cytoarchitecture and tissue microenvironment, and hence do not model the numerous intercellular interactions, proliferative behaviours and metabolic gradients characteristic of *in vivo* tissues ^4^. Critically, these features of *in vivo* tissues have been demonstrated to influence cell behaviour and responses to drug treatments ^5^. Therefore, utilising *in vitro* models that represent these crucial aspects of *in vivo* tissues is essential for reducing the failure rates of translating HTS data to clinical outcomes.

3D cell cultures represent critical features of *in vivo* tissues better than 2D cell cultures, meaning they may be able to more accurately predict therapeutic efficacy and drug responses ^6,7^. Such findings have significant implications not only for HTS, but fundamental biological research more generally. This understanding has facilitated a gradual movement towards the use of 3D cell cultures over 2D cell cultures for biological studies ^8^. However, the current lack of established workflows to produce large quantities of biologically relevant 3D cell cultures in an efficient and reproducible way, that are compatible with routinely used *in vitro* assays, remains a practical limitation to the widespread use of 3D cell culture models in HTS applications.

Many different methods have been developed over the past decades to create 3D cell cultures for biomedical research ^9^, but their technical limitations have largely restricted *in vitro* 3D cell culture to spheroid models, which have very little resemblance to the *in vivo* extracellular environment due to their free-floating nature ^10^. While spheroids enable a richer cell-to-cell interaction compared to 2D cultures, the highly influential role of the extracellular matrix in healthy and diseased tissue ^11^ is not addressed by this *in vitro* model option. Emerging 3D bioprinting technologies are addressing this shortcoming by generating tissue-like matrix-embedded 3D cell culture models at a quantity and consistency suitable for HTS in both small and large drug discovery projects.

RASTRUM™ is a drop-on-demand high throughput bioprinter that can deposit cells and matrix components into common tissue culture microplates to form 3D cell culture ^12^. High precision and delicate pressure regulation enables ejection of nanolitre volumes of different bioinks, which can be tuned to form hydrogels that match the mechanical and biochemical properties of different tissue types and provide a physiologically relevant matrix environment for cultured cells ^13^. After printing, further culture handling, such as media exchange, treatment addition and sample preparation, is identical to 2D cell culture and thus provides a cost-effective way for 3D cell culture generation and maintenance. The transparent, synthetic hydrogel matrices printed using RASTRUM™ do not interfere with standard imaging and assay techniques, and the microplate format provides compatibility of the 3D cell cultures with many common downstream analysis assays that have been developed for 2D cell cultures. In this way, RASTRUM™ provides a simple and automated workflow that efficiently generates reproducible matrix-embedded 3D cell culture models for the use in HTS.

Small and large molecule drug discovery projects, particularly high throughput screening, are widely using high sensitivity protein assays such as Alpha (amplified luminescence proximity homogeneous assay) ^14–16^. Alpha is a bead-based assay platform employing oxygen-channelling chemistry for the detection and quantification of various biomolecules, from small analytes such as cAMP and cytokines to intracellular and membrane bound protein complexes and biomolecular interactions^17^. The assays are characterised by no-wash mix-and-read protocol, high sensitivity, low sample requirements and easy automation, thus offer flexible solution for biomarker detection (AlphaLISA^®^) and intracellular kinase activity (AlphaLISA^®^ Surefire^®^ Ultra™) in cell based HTS assays.

The goal of this study was to establish a high throughput 3D cell culture workflow that allows the assessment of complex biological processes without complicating existing workflows. To achieve this, our workflow integrates RASTRUM™, a novel high throughput 3D bioprinting platform, and Alpha technology, a widely used HTS end point assay platform, to enable relevant intracellular and secreted molecule quantification in physiologically relevant 3D cell cultures (Figure 1). The workflow was assessed on breast cancer and fibroblast 3D cell cultures which were assayed for cytokine release and intracellular kinases. Conducting these experiments by combining the automated production of tissue-relevant 3D cell cultures with commonly used high throughput assays we aimed to lower the adoption hurdle for 3D cell cultures in productive HTS environments, especially for small molecule drug discovery workflows.

**Figure 1:**
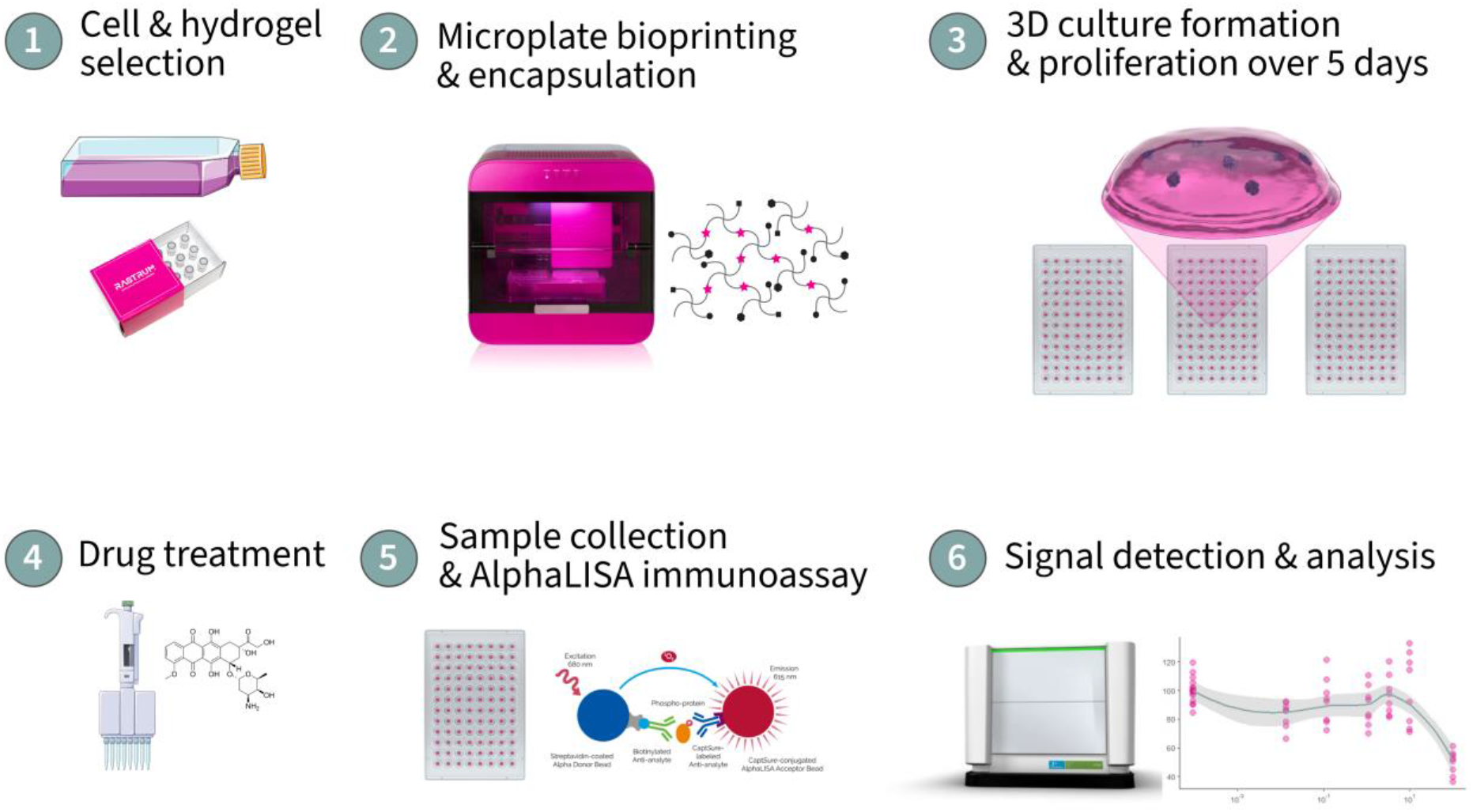
Workflow overview for high throughput target-based drug discovery with bioprinted 3D cell cultures; (1) cells are combined with their appropriate hydrogel formulation, (2) bioprinted with RASTRUM™ into microplates and encapsulated on impact inside the synthetic matrix. (3) 3D cell structures form during the proliferation and maintenance period until the treatment time point (4). Following the incubation with compounds of interest, the 3D cultures are processed with the standard AlphaLISA assay reagents (5) and quantified for intracellular or extracellular protein detection and analysis (6).

## Materials and Methods

### Cell Culture

#### Culture Details

**MCF-7 cells** (ATCC, HTB-22, obtained in 2014, human female adenocarcinoma of the mammary gland) and **MDA-MB-231 cells** (ATCC, HTB-26, obtained in 2014, human female adenocarcinoma of the mammary gland) were maintained in Dulbecco’s Modified Eagle Medium High Glucose (ThermoFisher Scientific, #11-965-118), 10% foetal bovine serum (Bovogen, #SFBS-F), and used between passage 8 and 17. **NHDF** (Sigma-Aldrich #106-05N, obtained in 2019, neonatal human dermal fibroblasts,) were maintained in Fibroblast Growth Medium (Sigma-Aldrich, #116-500), and used between passage 7 and 11. All cell cultures were maintained at 37 °C, 5% CO_2_, atmospheric O_2_ and 95% humidity without antibiotics in tissue culture-treated T75 flasks (Cellstar Greiner Bio One, Interpath Services #658175), and passaged at 80% confluence with Trypsin-EDTA (ThermoFisher Scientific #25300062).

#### 2D Culturing method

Cell cultures for 2D experiments were manually seeded with multichannel pipettes at 3500 (MCF-7 and MDA-MB-231) or 2000 (NHDF) cells per well into tissue culture treated 96 well plates (ThermoFisher Scientific #167008) in 150 μL culture media.

#### 3D Culturing method

3D hydrogel-encapsulated cell cultures for drug treatment experiments were bioprinted using a non-contact drop-on-demand 3D bioprinter (RASTRUM, Inventia Life Science). The bioprinter incorporates a flyby printhead which prints directly onto the substrate while in motion, leading to high-throughput printing of 3D cell cultures. Its 2-axis linear motion positioning system is capable of accurately positioning droplets on the substrate at a resolution of 20 μm along each axis. The droplet dispensing system consists of multiple independently addressable microvalves, which eject fluids at nanolitre precision. For the experiments in this study, we chose the 3D Small Plug Cell Model (dimensions: 2×2×0.5 mm), with polyethylene glycol (PEG) bioink formulations that mimic the mechanical and biochemical properties of the relevant *in vivo* tissue (Inventia Life Science, #Px01.00 for MCF-7, Px02.03P for MDA-MB-231, and Px01.02P for NHDF). The 3D cultures created by RASTRUM use two components: bioink and activator. Cell pellets are resuspended in activator solution which upon addition to the bioink by the bioprinter forms an instant gel. Each cell line was printed at 5×10^6^ cells/mL and loaded into the dedicated printer cartridge, alongside the bioink fluids. Following the automatic priming of all fluids into the nozzles, each cell line was printed into separate tissue culture plates, with Rastrum creating the 3D Small Plug across the 96 well plates within 10 minutes with an average of 700 cells per well (CV: 17%). Upon completion of the print run, 150 μL culture medium was added to each well via multichannel pipettes.

### Drug treatment

MCF-7 and MDA-MB-231 cultures were treated with doxorubicin (Sigma-Aldrich #44583, dissolved in DMSO), by replacing 50 μL cell culture medium with diluted doxorubicin to the relevant concentrations at no more than 0.1% v/v final volume. Control samples were treated with 0.1% DMSO (Sigma-Aldrich #D8418) diluted in culture medium. The penetration of doxorubicin was monitored by utilizing the autofluorescence of doxorubicin at 590 nm with an excitation at 480 nm, captured on an epifluorescence microscope (Zeiss Axio Observer 7).

NHDF cultures were treated with lipopolysaccharide (LPS, Sigma-Aldrich #L6529, dissolved in phosphate buffered saline), by replacing 50 μL cell culture medium with diluted LPS at the relevant concentrations.

### Intracellular and extracellular cell culture assays

#### Cell toxicity assays

Metabolic activity of 2D and 3D MDA-MB-231 cell cultures was assessed with CellTiter-Glo^®^ cell viability reagent (Promega, #G7570) according to the manufacturer’s instructions without any modification: the diluted CellTiter Glo substrate was added to the culture wells at the end of the treatment period, followed by a 10 min incubation, and luminescence quantification on a FLUOstar Optima (BMG labtech).

The Adenosine TriPhosphate (ATP)-monitoring luminescence assay ATPLite™ (PerkinElmer, #6016943) was used to quantitatively assess proliferation and cytotoxicity of 3D MCF-7 cultures. The ATPLite™ assay followed standard recommended no wash one step protocol and was read on an EnSight™ Multimode Plate Reader (PerkinElmer), without any deviation from the manufacturers protocol: 50 μL cell lysis solution was added to the 3D cell cultures and incubated for 5 minutes on an orbital shaker, followed by the addition of 50 μL ATPLite substrate solution and a further 5-minute shaking. The culture plates were then incubated for 10 minutes in the dark, before the quantification of their luminescence intensity.

#### Intracellular kinase activity

AlphaLISA® SureFire® Ultra™ cellular kinase immunoassays were used to measure endogenous kinase activity of p-mTOR (Ser2448) (PerkinElmer, #ALSU-PMTOR-C500), p-ERK 1/2 (Thr202/Tyr204) (PerkinElmer, #ALSU-PERK-A500), p-GSK-3β (Ser9) (PerkinElmer, #ALSU-PGS3B-A500). Additionally, ERK 1/2 Total (PerkinElmer, #ALSU-TERK-A500) and Cofilin Total (PerkinElmer, #ALSU-TCOF-A500) were used as a normalisation for phosphorylation studies. For each study, all targets were assayed using the same cell culture lysates from single 2D or 3D cultures, enabling the measurement of multiple kinase activity levels in the same cell culture sample. The AlphaLISA® SureFire® Ultra™ assay was run in total volume 20 μL (using only 10 μL cell culture lysate sample) in 384 well plate (PerkinElmer, #6007290) following standard assay two step no-wash protocol. We modified the manufacturers protocol of this assay to increase the efficient extraction of intracellular proteins from the embedded 3D cultures. The incubation period of the lysis buffer (70 μL per well) was extended to 40 min, with five times pipetting up and down at 15 and 30 min incubation. The complete lysate was then sampled for further processing in line with the manufacturers protocol in 384 well plates. After the lysis step, the cell constructs (spheroids or networks) were largely destroyed and cell lysis was visually confirmed, with near empty gel left behind in each well. The assay was read on an EnSight™ Multimode Plate Reader (PerkinElmer).

#### Cytokine release assay

AlphaLISA™ Immunoassay was used to assess the IL-6 cytokine release (PerkinElmer, #AL223) in the media in 2D and 3D fibroblast cell cultures. The AlphaLISA™ assay was in 20 μL reactions (requiring only 2 μL cell culture media sample) in 384-well plate (6007290, PerkinElmer) following standard two-step no-wash protocol without any modifications. The assay was read on an EnSight™ Multimode Plate Reader (PerkinElmer).

### Statistical Analysis

Data analyses were performed with R in RStudio Team (2020) (RRID:SCR_000432, RStudio: Integrated Development for R. RStudio, PBC, Boston, MA, USA). Treatment effects were assessed using one-way or two-way analysis of variance (ANOVA) as relevant, followed by Bonferroni’s multiple comparisons test where appropriate. Experiments were conducted with at least three biological replicates and at least two technical replicates. Significance was accepted at p<0.05 and data presented as mean ± standard error of mean (SEM) for biological replicates.

## Results

The goal of this protocol paper was to establish a workflow that would generate tissue-relevant 3D cell culture models which are compatible with widely used in vitro assays at a scale suitable for HTS projects. We conducted three different validation experiments to assess the utility of this workflow by 1) comparing the treatment responses between 3D cultures and 2D cultures of the same cell type; 2) monitoring the penetration of an anti-cancer drug into 3D cultures and its effect on intracellular kinase activity; and 3) determining the reliable detection of released factors from matrix-embedded cells.

### 3D and 2D MDA-MB-231 cultures were equally susceptible to doxorubicin treatment

MDA-MB-231 cells exhibit a spindle-like, network-forming morphology in 3D, and similarly develop elongated forms in 2D, representative of their metastatic cancer source ^18^. MDA cells readily proliferated and migrated through the RGD-containing hydrogel, with a mature morphology exhibited after four days in culture. The general drug responsiveness to DOX was visible within 48 h for both 2D and 3D cultures (Figure 2A and Figure 2B respectively) and confirmed via CellTiterGlo (Figure 2C), showing a near-identical iC50 value between both culture environments (2D: 2.1 μM, 3D: 2.9 μM, Figure 2C), which is in line with previous reports ^6,7,19^.

**Figure 2:**
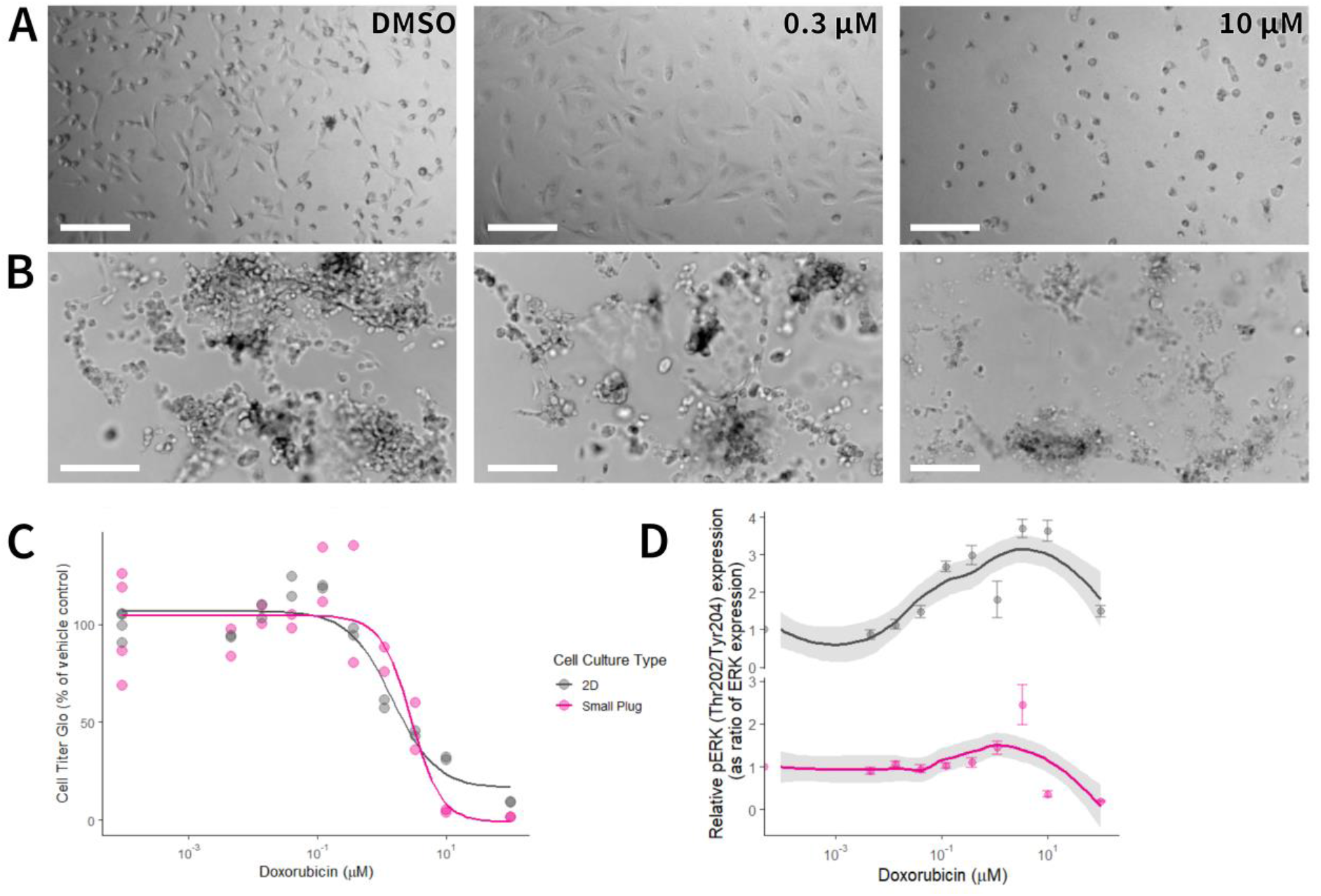
Doxorubicin reduces viability and increases ERK phosphorylation in 2D and 3D MDA-MB-231 cultures. Representative images of 2D (A) and 3D (B) MDA-MB-231 cultures show the doxorubicin treatment response after 24 h, scale bar = 200 μm. Viability of treated cultures was assessed after 24 h via Cell Titre Glo (C, data represent individual biological repeats), and phosphorylation of ERK as ratio to total ERK quantified via AlphaLISA from intact cultures (D, n = 3 per concentration and culture type, data represent mean ± SEM).

While viability assays are widely used in 2D and spheroid 3D cultures, assessing treatment effects on intracellular protein kinases has become a core component of advanced screens of novel compounds. The phosphorylation status of ERK has been implicated in both cancer proliferation and anti-cancer drug response ^20^. We therefore quantified intracellular ERK 1/2 phosphorylation at Thr202/Tyr204 and identified a concentration-dependent phosphorylation increase in 2D cultures and decrease in 3D cultures, relative to total ERK signal (Figure 2D). In 2D cultures, the pERK/ERK ratio increase peaked at 3.33 μM and 10 μM (3.69±0.25, 3.63±0.28 respectively), while in 3D cultures 10 μM and 100 μM doxorubicin resulted in a reduced phosporylation ratio (0.36±0.06, 0.19±0.019 respectively).

### Doxorubicin readily enters embedded MCF-7 spheroids and alters intracellular GSK3β phosphorylation

Published studies show that doxorubicin treatment effectively leads to the death of MCF-7 breast cancer cells in 2D and at higher concentrations in 3D spheroid cultures ^19^. We replicated these experiments to monitor both drug penetration into hydrogel-embedded MCF-7 spheroids, and ease of analysing intracellular kinase response to the drug treatment. MCF-7 spheroids (average surface area: 1480 μm^2^) showed substantial morphological changes after 24 h of doxorubicin treatment at and above 10 μM, with no intact spheroids present at 100 μM (Figure 3A). Fluorescent microscopy imaging confirmed the penetration of doxorubicin into the MCF-7 spheroid (Figure 3B). Measuring intracellular ATP quantities with ATPLite showed a marked increase in ATP at 3.33 μM and 10 μM (Figure 3C), likely due to increased autophagy activity^21^. We were able to readily sample the intracellular proteins of the hydrogel cultures with the regular AlphaLISA kit reagents and upon quantifying the phosphorylation of GSK3β, detected a market decrease in Ser9 phosphorylation relative to our loading control (Cofilin) at 100 μM doxorubicin (49.2% ±3.67, Figure 3D).

**Figure 3:**
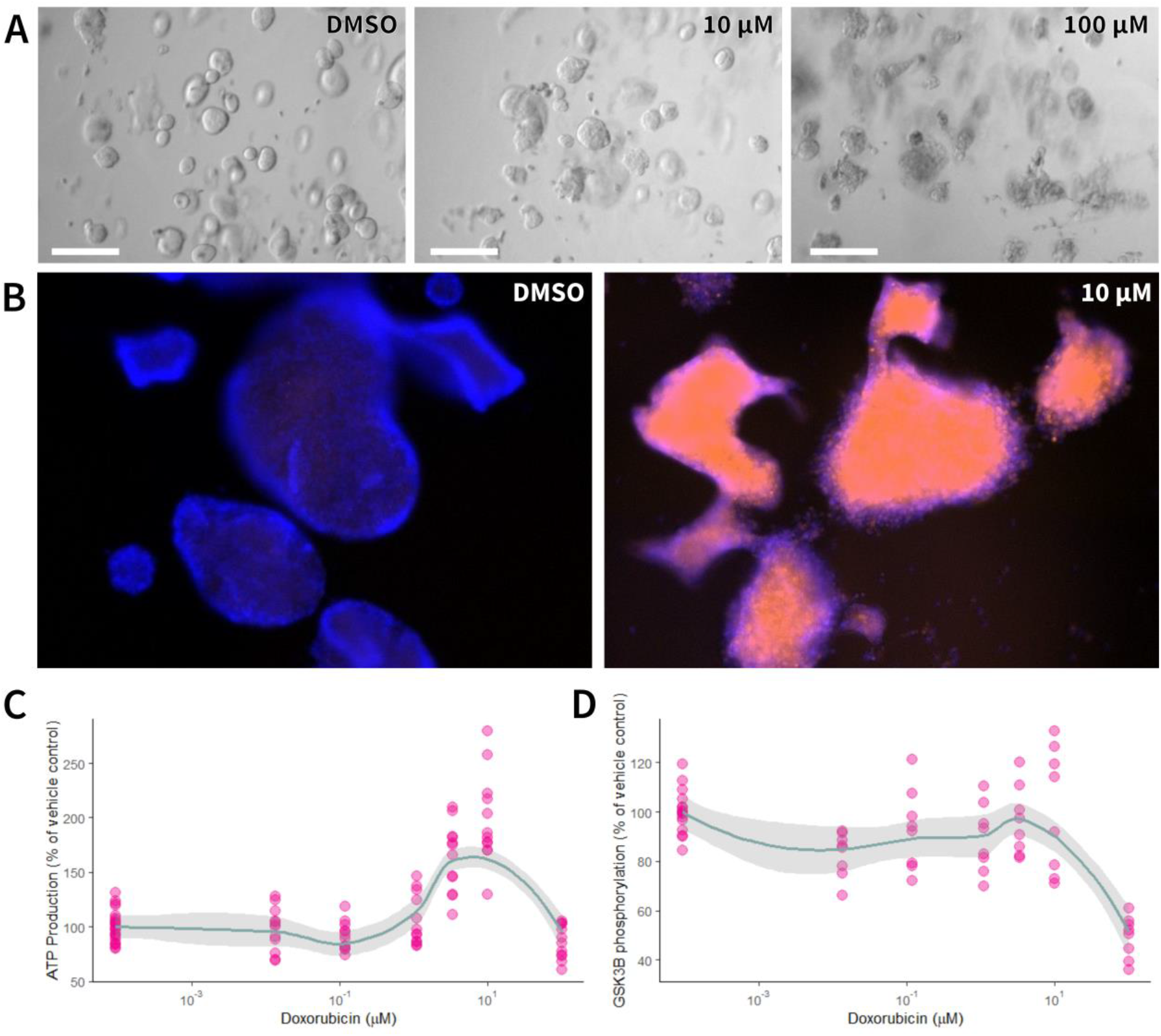
Doxorubicin readily penetrates 3D MCF-7 cultures, resulting in a dose-dependent ATP and GSK3β phosphorylation change. Embedded MCF-7 cultures were treated with DOX or DMSO vehicle for 24 h (A, scale bar = 200 μm), with its penetration confirmed by monitoring the autofluorescence of DOX in untreated (B) and treated spheroid cultures. The treatment effect on intracellular ATP concentration (C) and GSK3β phosphorylation (D) was assessed after 24 h. Data normalised to untreated samples, with each data point representing individual cultures, mean line and shaded standard error of mean.

### Cytokine release patterns can readily be detected in 3D hydrogel fibroblast cultures

The possibility of creating consistent 3D cell cultures at quantities suitable for HTS also opens the potential for tissue-relevant co-culture models. Recently, pro-inflammatory signaling by fibroblasts has moved into the focus of cancer treatment research for its contribution to cancer drug resistance ^22^. We therefore created 3D dermal fibroblast cultures in our third workflow validation experiment to assess the release of the pro-inflammatory cytokine interleukin 6 (IL-6) following stimulation with bacteria wall segments (lipopolysaccharide, LPS) and compare the activation profile with 2D cultures. Fibroblasts formed extensive networks in 3D printed hydrogel cultures (Figure 4A), substantially more complex than 2D cultures of the same age (Figure 4B). Treatment with LPS over 48 h triggered a concentration-dependent release of IL-6 into the culture media, which was readily detectable via the AlphaLISA assay in both 2D and 3D cultures (Figure 4C). The Ec50 LPS-induced IL-6 release was near identical to both culture types (2D: 0.06 μg/ml, 3D: 0.04 μg/ml), with a larger effect size in 2D cultures than 3D cultures (1500% and 200% respectively, Figure 5c). The near identical response pattern of both culture types suggests not just an unhindered penetration of LPS into the hydrogel, but also the diffusion of IL-6 from the cells into the general culture media.

**Figure 4:**
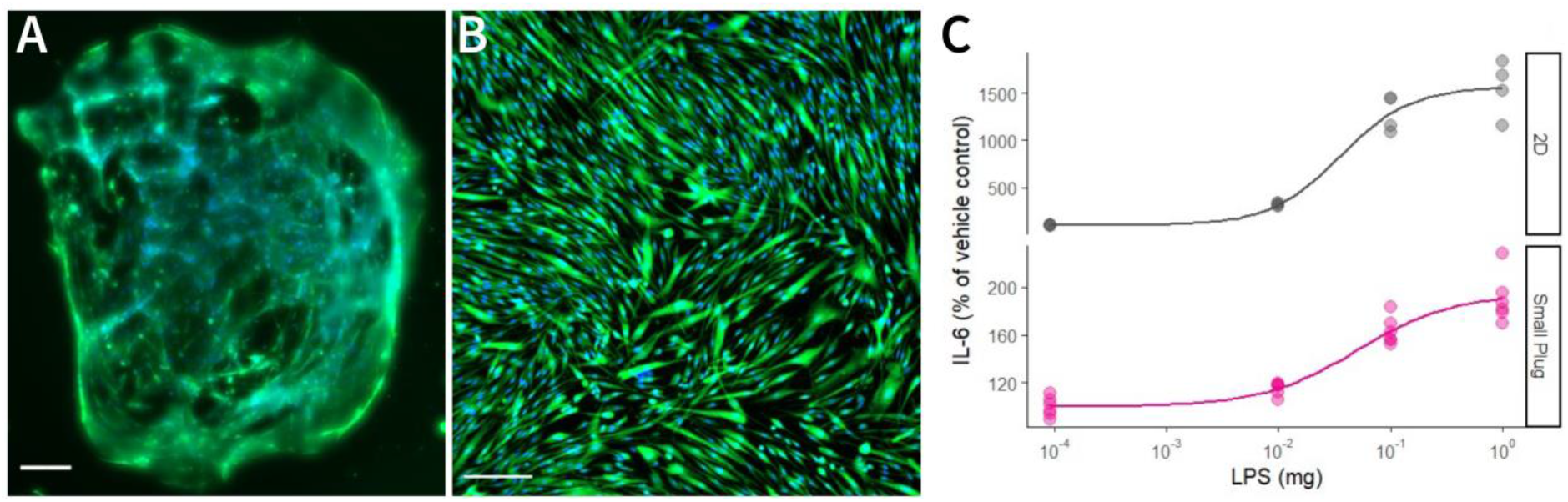
Release of pro-inflammatory cytokines by stimulated fibroblasts was readily detected in both 3D and 2D cultures. Five days old 3D (A) and 2D (B) dermal fibroblasts were treated with LPS for 48 h, after which extracellular IL-6 presence in the culture media was quantified via ALPHALisa (C). Data normalised to untreated samples, with each data point representing individual cultures (scale bar = 200 μm).

## Discussion & Conclusion

End point assays are standard experiments in small molecule cell-based screening, however they are less commonly used in 3D cell culture based screens due to challenges of tissue-relevant 3D cell culture generation, handling, and compatibility with screening assays. The workflow proposed here offers an alternative to the traditional 2D small molecule drug screening workflow, utilising end point assay as a final readout, with the only difference being the use of 3D cell culture as a starting point.

The workflow was established and validated with breast cancer and fibroblast models, by performing already published 2D cell culture experiments for cytokine release and intracellular kinase response to small molecule stimulation.

The RASTRUM 3D bioprinter offered an easy and automated printing workflow that generated reproducible hydrogel-embedded 3D cell culture models. Once the culture were printed, further handling such as media exchange, culture treatments and sample preparation were identical to 2D cell culture and thus offers cost effective and efficient 3D cell culture generation. The hydrogel matrix used in the printing method did not interfere with the standard imaging and assay techniques, while making fluid handling easier than with free-floating 3D spheroid cultures.

Our results show that the RASTRUM hydrogel 3D tissue cultures can readily be used with several standard 2D HTS assays, and more importantly with advanced protein assays, enabling multi-target detection and thus maximising the output of each culture. The doxorubicin and LPS treatment responses in our reference experiments are in line with published findings from regular 2D and 3D spheroid assays.

Further, handling time of cultures and assays was identical between 2D and 3D cultures and both delivered enough material for Alpha assay multitarget detection as shown with both the MDA-MB-231 and MCF-7 experiments. This ease of use is relevant for the adoption of novel sample sources; however the true value of our approach is derived by using more physiologically relevant in vitro models to study cell biology and cell response to the drug stimulation compared to both 2D and spheroid cultures ^23^. Tailoring the hydrogel matrix to each tissue niche offers closer prediction of the in vivo response to treatments than 2D cultures would, and consequently a more wholistic insight into the biological response of the cell ^11^. While our workflow focuses on the Alpha assays to quantify specific proteins, the wide variety of available formats and the high customisability of this technology makes this assay choice nonetheless relevant to many high throughput experimental settings in drug discovery using both 2D and 3D cell cultures.

Notwithstanding our use of 96-well cell culture formats in this manuscript, the presented workflow can directly be used in 384 well plates, as all components are available for this format. Taking this aim further, the next steps will be the full incorporation of the currently standalone 3D bioprinting into a truly high throughput setting, by automating the entire workflow from the starting cell culture, over 3D printing and culture handling, to assay processing that is achieved by fully integrated automation. Such a walk-away workflow is comparable to many current 2D cell culture screening approaches in a HTS environment and will bring the often-described potential of advanced 3D tissue culture for medical discoveries to fruition in translation-focused research settings.

## Author contributions

M.E. and M.S. conceived the project, and coordinated scientific work; M.E., L.B. and B.A. designed and performed the cell culture experiments and performed data analysis; M.E., L.B. B.A. and M.S. wrote the paper.

